# Pro-restitutive *Bacteroides thetaiotaomicron* reprograms the transcriptome of intestinal epithelial cells by modulating the expression of genes essential for proliferation and migration

**DOI:** 10.1101/2025.09.30.679439

**Authors:** Angela Gao, Violet Newhart, Madison Flory, Ashfaqul Alam

## Abstract

The mammalian intestine harbors a highly complex, very diverse, and numerically vast community of symbiotic microorganisms, which profoundly influence the development and maintenance of the intestinal barrier function. Alterations in microbial composition, known as dysbiosis, are observed in Inflammatory Bowel Disease (IBD), colorectal cancer (CRC), and gastrointestinal infections; however, the exact causal relationship between these changes and the resolution of intestinal inflammation and the repair of damaged mucosa remains unclear. Notably, IBD is not only marked by dysbiosis but also by changes in microbial metabolic pathways and metabolite landscape in the intestinal lumen. The small molecules and microbial metabolites present in the intestinal lumen have emerged as potential regulators of gut pathology, cancer, and mucosal repair. Investigating how altered microbiota and microbial metabolic activities influence intestinal epithelial cells (IEC) can provide insights into their role in the regeneration of mucosal epithelia and restoration of gut barrier functions. This knowledge can be harnessed to promote intestinal homeostasis, prevent relapse, and prolong remission of IBD. To dissect the complex interplay between the gut microbiome and IEC, we focused on the overrepresented bacterium *Bacteroides thetaiotaomicron*. Here, we show that *B. thetaiotaomicron and Akkermansia muciniphila*, the dominant members of gut microbiota, expand during the repair & resolution phase of the chemically induced acute murine colitis. Furthermore, our bioinformatics analysis demonstrated that the elevated relative abundance of *B. thetaiotamicron* was also accompanied by rewiring of bacterial metabolic programs towards the essential amino acid metabolism, polyamine synthesis and utilization, stress response mechanisms, cell envelope biogenesis, and nutrient scavenging. Our RNA sequencing and transcriptomic analysis of primary human colonic epithelial cells cocultured with *B. thetaiotaomicron* showed that *B. thetaiotaomicron* stimulates the expression of genes and pathways involved in different cellular functions, including proliferation, differentiation, adhesion, lipid metabolism, migration, chemotaxis, and receptor expression. Our study emphasizes the crucial functions of the gut microbiome and metabolic activities in regulating the functions of intestinal epithelial cells during the repair of injured gut mucosa. Thus, these microorganisms and their metabolism hold promise as potential therapeutic agents.

## Introduction

The mammalian gastrointestinal tract harbors a highly complex, very diverse, and numerically vast community of symbiotic microorganisms, including bacteria, viruses, protozoans, and fungi. The composition and function of the gut microbiome are significantly influenced by birth conditions, geographic location, dietary habits, infectious diseases, medical interventions, and other socio-biological factors^1-5^. Alterations in the microbiome across the lifespan can substantially affect human health, as the microbiome modulates vitamin synthesis, nutrient digestion, drug metabolism, immune responses, angiogenesis, bone density, and neural function. Given its direct interaction with the gastrointestinal tract, the altered gut microbiome, often termed “dysbiotic” microbiota, plays a critical role in the development of intestinal inflammation^10^.

Although numerous factors contribute to the development and progression of inflammatory bowel disease (IBD), the gut microbiome plays a pivotal role in this multifaceted interplay^1-5^. The interaction between the microbiome and the intestine is complex and dynamic, making it a vital area of research as we aim to understand the etiology and consequences of IBD^1-7^.

IBD encompasses both ulcerative colitis (UC) and Crohn’s disease (CD). Although both CD and UC share some similar clinical manifestations and symptoms, each exhibits distinct pathological features. In UC, inflammation is restricted to the colon, specifically the mucosal layer of the colon ^2^. Inflamed regions can be isolated, located among healthy parts of the colon. Conversely, in CD, inflammation may occur anywhere along the gastrointestinal tract and can affect deeper layers of the intestines^8,9^.

IBD is characterized by cyclical patterns of disease activity. During remission, patients usually experience few symptoms, reflecting a temporary resolution of inflammation. At other times, patients face exacerbations, inflammation, or flares, during which clinical symptoms recur and may eventually lead to intestinal damage. Although there is no cure for IBD, current treatments aim to control and manage IBD flares, including steroids, anti-inflammatory drugs, and biologic therapies such as monoclonal antibodies, all designed to attenuate inflammation^8,9^. The longer a person stays in a state of active inflammation, the higher their risk of developing later complications, including intestinal infections and colitis-associated colorectal cancer (CAC). Maintaining remission is a complex process influenced by a combination of medical, dietary, environmental, and psychosocial factors^7-10^. The gut microbiota profoundly affects remission by influencing immune activation, intestinal barrier integrity, and producing beneficial metabolites. It has been previously reported that the gut microbiota of UC patients who achieve long-term remission shows evidence of significant recovery of the gut microbial ecosystem, with diversity and composition resembling that of healthy gut microbiota, and also displaying an abundance dominated by *Bacteroides thetaiotaomicron* or *Akkermansia muciniphila* (Claudia et al., 2023).

Although the gut microbiota has been extensively investigated for its role in modulating inflammatory processes, and numerous associations have been identified, the precise roles and metabolic contributions of many individual microbes to the remission of IBD and regeneration of injured intestinal mucosa are not yet fully understood^7-15^. The four predominant phyla in the gut microbiome—Firmicutes, Bacteroides, Proteobacteria, and Actinobacteria—constitute the majority of microbial populations, with additional minor representation from other phyla. In IBD, reductions in Firmicutes and increases in Proteobacteria are frequently observed. However, it remains to be elucidated whether these microbiota alterations, especially those involving the Bacteroides genus, actively participate in the pathogenesis or resolution of intestinal inflammation. IBD is characterized by both alterations in the microbes themselves and changes in their downstream metabolic pathways and metabolic products^10-15^. These microbial metabolites can influence various physiological aspects of the IECs and can thus affect IBD in either a protective or detrimental manner. Furthermore, these microbial compositions and metabolic shifts often lead to reprogramming of the IECs’ metabolic functions, including energy metabolism, amino acid catabolism, biosynthetic pathways, bile acid transformation, and signaling pathways. Thus, dysbiosis in the microbial community or its altered metabolic landscape may result in compromised intestinal barrier function, increased mucosal inflammation, impaired mucosal healing, and the negative outcome of IBD^14,15^.

The regenerative capacity of the intestinal epithelium following injury is governed by finely tuned complex processes of cellular proliferation and migration emanating from crypt compartments adjacent to the wounded mucosa^1,41,54^. The maintenance of epithelial integrity in the gut is fundamentally dependent on robust proliferative responses, especially in the aftermath of injury. We have previously demonstrated that commensals such as *Lactobacillus rhamnosus* GG engage in the repair of both mechanical and chemical injuries in murine models. The administration of *L. rhamnosus* GG and *A. muciniphila* significantly accelerates wound closure and crypt regeneration through a redox-dependent signaling pathway—an essential prerequisite for the downstream activation of focal adhesion kinase (FAK) and ERK in enterocytes of the mouse’s distal colon^1,41,54^.

*Bacteroides thetaiotaomicron* is a keystone commensal bacterium in the human gut, known for its multifaceted contributions to nutrient assimilation and immune modulation. Seminal work by Hooper et al. (2001) revealed that *B. thetaiotaomicron* stimulates the expression of angiogenin-4—a bactericidal protein secreted by Paneth cells—thereby influencing both microbial ecology and host defense mechanisms^3^. Moreover, *B. thetaiotaomicron* regulates epithelial differentiation and fine-tunes nutrient transporter expression, collectively advancing epithelial homeostasis. Thus, *B. thetaiotaomicron* emerged as a central architect in the maintenance and adaptation of host intestinal physiology. Interestingly, *B. thetaiotaomicron* has been shown to promote the post-antibiotic expansion of *Salmonella typhimurium* by catabolizing sialic acid with sialidase^56^. In contrast, *B. thetaiotaomicron* has been reported to produce propionic acid, which in turn helps protect against certain enteric infections. *B. thetaiotaomicron* is a key gut microbiota with a wide range of enzymatic activities to scavenge carbohydrates and proteins, which could significantly influence the health and stability of the intestinal mucosa by modulating the outcome of enteric infections as well as IEC function. Therefore, additional research is necessary to understand the complex roles of *B. thetaiotaomicron* in regulating intestinal barrier integrity.

To understand the effects of the microbiome and its metabolites on colitis and mucosal repair, we identified an overrepresented bacterial species – *B. thetaiotaomicron* — and its impact on cultured IECs^42,44,45^. Here, we demonstrate that *B. thetaiotaomicron* and *Akkermansia muciniphila*, the dominant gut microbiota members, expand during the repair and resolution phase of chemically induced acute murine colitis. Additionally, our bioinformatics analysis revealed that the increased relative abundance of *B. thetaiotaomicron* was accompanied by reprogramming of bacterial metabolic pathways toward essential amino acid metabolism, polyamine synthesis and utilization, stress response mechanisms, cell envelope biogenesis, and nutrient scavenging. Our RNA sequencing and transcriptomic analysis of cultured colonic epithelial cells cocultured with *B. thetaiotaomicron* indicated that this bacterium stimulates gene and pathway expression involved in various cellular functions, including proliferation, differentiation, adhesion, lipid metabolism, migration, chemotaxis, and receptor expression. Thus, *B. thetaiotaomicron*, which is enriched during the recovery phase of murine colitis, promotes genes involved in proliferation, migration, and metabolic rewiring, indicating a potential role of B. *thetaiotaomicron* in mucosal wound repair and homeostasis.

## Results

### Alterations in colonic gut microbiota during the recovery phase of chemically induced murine colitis

We previously demonstrated that the microenvironment of injured murine gut alters a local pro-restitutive microbiota^1^. However, we do not completely understand how the acute inflammation stimulates alterations in microbiota, which affect the gene expression of intestinal epithelial cells during restitution. The DSS colitis mouse model is widely used to investigate the interplay between intestinal inflammation and microbial communities. Yet, the specific changes in microbial communities and their metabolic functions during the healing and regeneration stages of DSS colitis are not well understood. Therefore, to model acute intestinal inflammation followed by mucosal repair, mice were given 4% DSS for 7 days, followed by 7 days of water to allow repair of the injured intestinal mucosa. We collected stools on day 14 (1-week post DSS-removal; Figure 1A). DNA was purified from the stools as well as from adjacent intact mucosa and local luminal contents. Bacterial 16S rRNA genes (V4 region) were PCR-amplified, and the amplicons were sequenced by a high-throughput sequencing (HTS) platform. At the phyla level, microbiota analysis revealed that during the recovery phase after DSS treatment, stool microbiota consisted of a lower abundance of Bacillota and a higher abundance of Pseudomonadota, Verrucomicrobiota and Bacteroidota compared to the microbiota of the control mice. Furthermore, we determined the alpha diversity indices (Figure 1B). The Shannon index indicated a decrease in microbial taxonomic diversity during the recovery phase compared to the control mice. Similarly, the Simpson index revealed that there are significant differences in microbiome evenness 7 days after DSS treatment. In addition, we found significantly increased abundances of *Ruminococcus (genus), Akkermansia (genus), Fusobacteriota (phylum), Escherichia (genus), Prevotella (genus), Bukhoderia (genus)*, and *Bacteroides thetaiotamicron* (Figure 1A-C). In contrast, at the genus level, the control group has higher levels of *Acetohalobium, Anaerostipes, Desulfovibrio, Fibrobacter, Lacrimispora, Ligilactobacillus*, and *Megalobacillus*. Figure 2C shows a heatmap illustrating the depletion of six genera in the 7-day post-DSS treatment compared to the control mice. The initial increase in Bacteroidota phyla prompted a further investigation into the enrichment of specific bacterial species in these models. Thus, we performed enrichment analysis at the genus level, which showed enrichment of *B. thetaiotaomicron* at 7 days post-DSS treatment. The other enriched species (Figure 2B) in the recovery phase include *Barnesiella viscericola, Paludibacter propionicigenes, Akkermansia muciniphila, Lachnospirigenes, Basilea psittacipulmonis, Brevundimonas subvibrioides, Escherichia coli, Lactobacillus kefiranofaciens, Blautia obeum*, and *Ruminococcus champanellensis*. On the other hand, the species that were depleted in the recovery phase include *Anaerostipes hadrus, Thermanaerobacter marianensis, Lachnispira saccharolytica, Coprococcus catus, Agathobacter rectalis, Lachnoclostridium phytofermentans, Ruminococcus sp. SR1/5, Alistipes finegoldii, Butyrivibrio proteoclasticus, Ruminococcus torques, Acetobacterium woodie*, and *Desulfosporosinus meridiei*.

**Figure 1.**
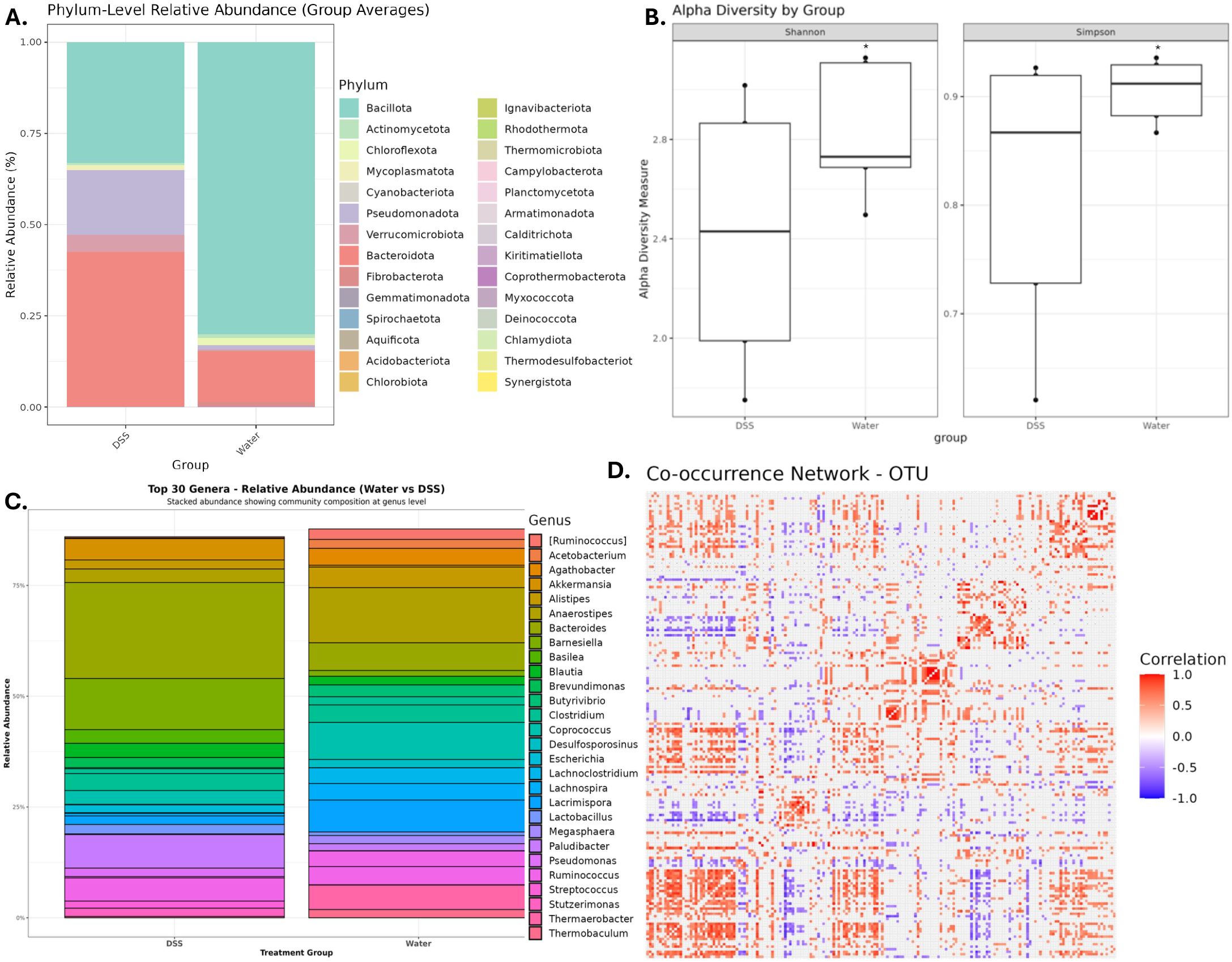
Microbiome profiling reveals characteristic changes in murine models of chemically induced colitis during the recovery phase of the disease model, 7 days after DSS withdrawal. Mice were subjected to 4% DSS colitis for 7 days, followed by recovery from colitis for 7 days with water. (A) Mean relative abundance of bacterial phyla determined by 16s rRNA gene sequencing and microbiome analysis. (B) Alpha diversity estimates (Shannon on the left; Simpson on the right) show a positive correlation between reduced bacterial taxa diversity and DSS colitis. (C) Specific changes in bacterial genus at 7 days post-DSS treatment. (D) Analysis of the co-occurrence network of microbial taxa in the luminal contents of mice during recovery, 7 days after DSS treatment. N = 5 per group. *P < 0.05; Student’s t-test.

**Figure 2.**
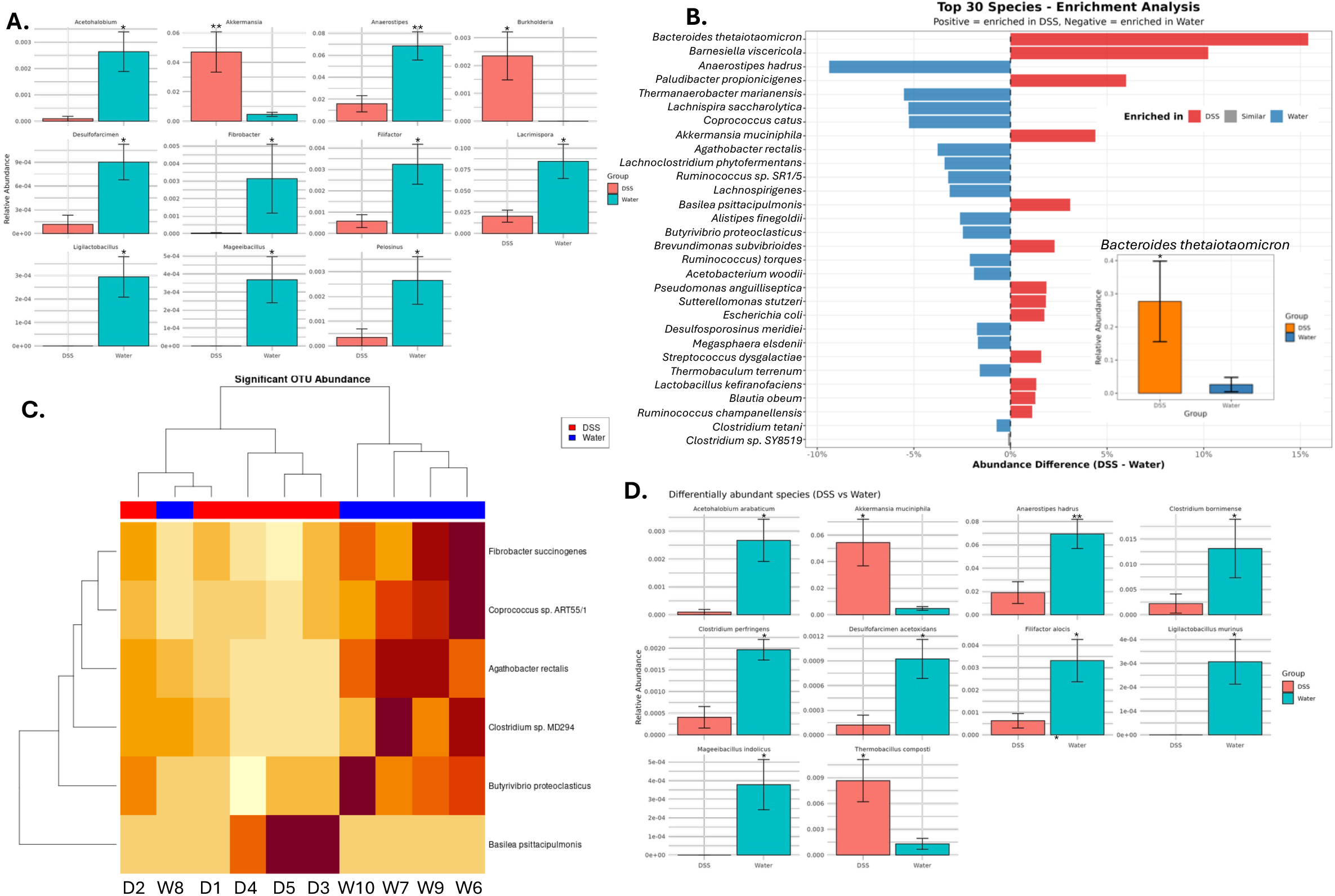
Microbiome profiling reveals alterations in the relative abundance of colonic bacterial genus and species during the recovery phase of chemically induced colitis in mice, 7 days after DSS withdrawal. (A) Mean relative abundance of bacterial genus determined by 16s rRNA gene sequencing and microbiome analysis. (B) Enrichment analysis of the top 30 bacterial species. Inset, specific increases of *B. thetaiotamicron* during the recovery phase, 7 days post DSS withdrawal. (C) Heatmap shows dramatic alterations of 6 bacterial OTU abundance in response to DSS treatment. (D) Mean relative abundance of bacterial species determined by 16s rRNA gene sequencing and microbiome analysis. N = 5 per group. *P < 0.05, **P < 0.01; Student’s t-test.

Gram-negative species listed (Figure 2B-D) include Bacteroides thetaiotaomicron, *Barnesiella viscericola, Paludibacter propionicigenes, Akkermansia muciniphila, Basilea psittacipulmonis, Brevundimonas subvibrioides*, and *Escherichia coli*. Gram-positive members include *Lachnospiraceae (Lachnospirigenes), Blautia obeum, Ruminococcus champanellensis*, and *Lactobacillus kefiranofaciens*. This division reflects fundamental differences in cell envelope structure that influence interactions with the host, susceptibility to antibiotics, and mechanisms of secretion. Additionally, most of the top symbionts identified during this stage of recovery phase are anaerobic bacteria adapted to the anoxic colonic lumen. However, facultatively anaerobic *E. coli*, aerotolerant Lactobacillus species, and also *Brevundimonas* and *Basilea*—more oxygen-tolerant and not typical strict gut anaerobes—are also significantly increased. Thus, these data demonstrate that microbial composition profoundly alters during the recovery phase of chemically induced murine colitis.

Human IBD studies often demonstrate changes in microbial community composition characterized by reduced diversity, decreased anaerobic Firmicutes, and lower levels of mucin-utilizer *A. muciniphila* and *B. thetaiotamicron*, with elevated levels of *E. coli* and Enterobacteriaceae. We also observed an increase in *E. coli* levels; however, unlike previous human IBD studies, we found higher levels of *B. thetaiotamicron* and *A. muciniphila* during the recovery phase of the 7-day post-DSS withdrawal. Thus, our microbiome analysis revealed a unique signature in the bacterial community during the recovery phase post-acute colitis in the mouse intestine.

### Alterations in colonic gut microbiota are accompanied by predicted metabolic functions during the recovery phase of chemically induced murine colitis

After determining the profound alterations in microbial profiles, we next determined whether the microbial dysbiosis is also associated with the functional impairment in the microbial community’s metabolic capacity. To predict these metabolic changes from the 16s rRNA sequencing data, we performed a phylogenetic investigation of the communities by reconstructing unobserved states. This tool compared the predicted metagenomes between control mice (water) and mice recovering from DSS-induced colitis. As shown in Figure 3A, we observed a dramatically altered functional and metabolic potential of the gut microbiota compared to the control groups. During the 7-day post-DSS withdrawal, there was a significantly reprogrammed microbial metabolism involving a shift in many biological pathways. As shown in Figure 3B-D, we determined specific KEGG pathways that significantly increased in the gut microbial community 7 days post DSS withdrawal compared to the control group administered with regular water. We found an increase in microbial metabolism of L-arginine degradation and L-ornithine degradation pathways. These pathways are involved in the bacterial biosynthesis of putrescine, spermine, and spermidine, which are important for maintaining intestinal epithelial integrity. Furthermore, the concurrent upregulation of glycolysis and methylglyoxal degradation suggests a state of high, inefficient fermentative activity. Additionally, there is a trigger for LPS biosynthesis (Figure 3B-C). Moreover, there was an increase in amino acid metabolism and purine nucleotide biosynthesis pathways. Pathways such as ergothioneine biosynthesis (a potent antioxidant) and heme b biosynthesis indicate that the bacteria are indeed in need of metabolic reprogramming in a stressed environment caused by inflammation, antibacterial peptides, reactive oxygen species, and similar host responses (Figure 3B-D).

**Figure 3.**
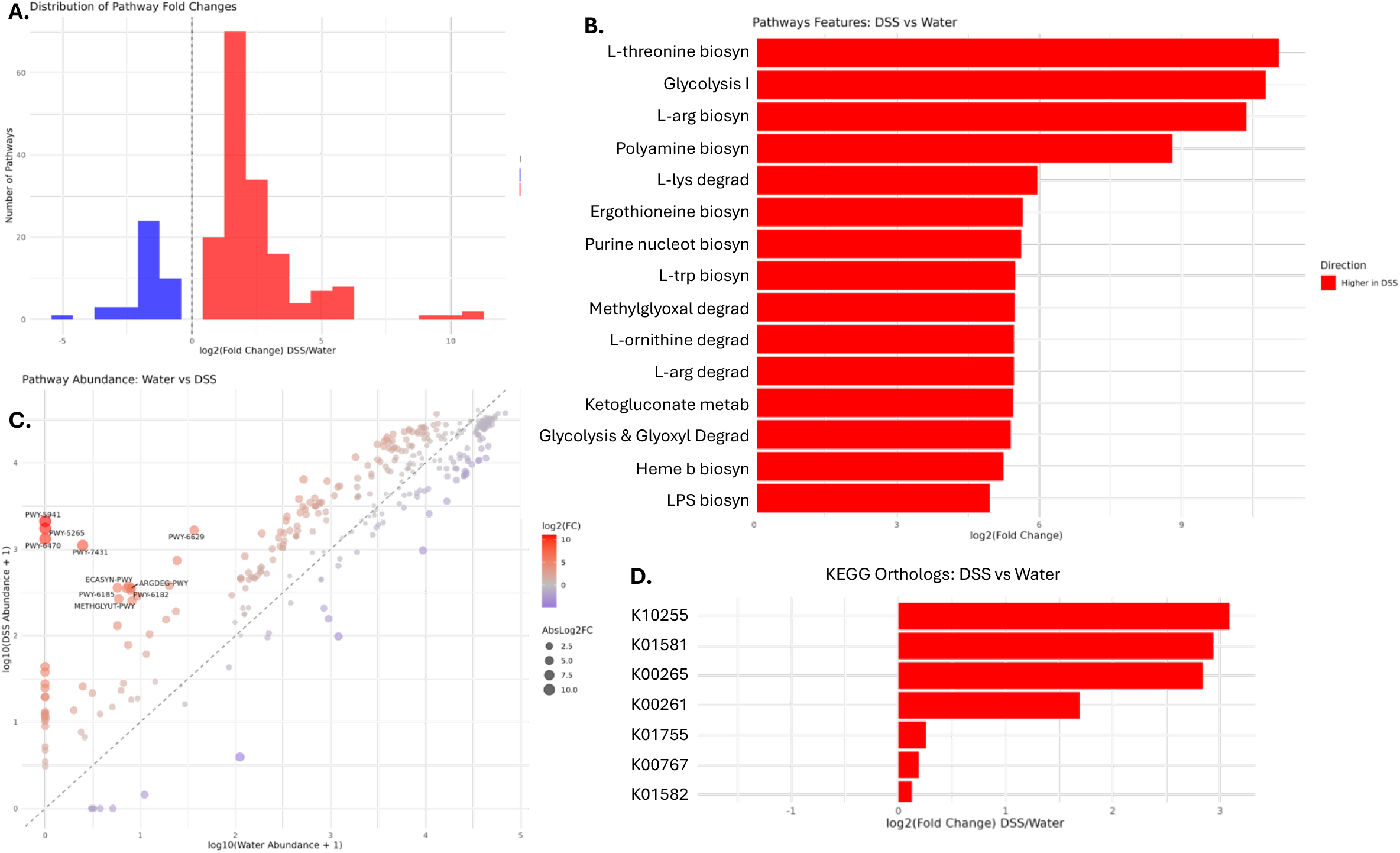
Bioinformatics analysis of bacterial KEGG Orthologies (KO) prediction and pathway enrichment in the lumen during the recovery phase of chemically induced colitis in mice, 7 days post DSS withdrawal. (A) The distribution of pathway fold changes in two different conditions (blue is higher in control mice with water; red is higher in mice treated with DSS). (B) KEGG Pathway Enrichment analysis of bacterial metabolic pathways altered in the luminal contents of mice recovering from DSS-induced colitis. (C) The scatter plot displays the predicted microbial pathway abundances, derived from 16S rRNA gene sequencing-based taxonomic profiling and PICRUSt2 analysis. Each data point corresponds to a specific KEGG or MetaCyc metabolic pathway, providing insights into functional potential within the microbiome. *P < 0.05

Interestingly, Bacteroides species are known to be producers of polyamine metabolites, indicating that we would expect an increase in polyamines in the acute colitis models due to the rise in Bacteroides species. Previously, we showed in a mouse model that when germ-free mice were colonized with *B. uniformis*, spermidine levels increased in the intestinal content, suggesting that gut bacteria, especially those associated with IBD, contribute to excess polyamine production. These consistent increases in specific bacterial groups, such as Bacteroidota phyla and the highly enriched *B. thetaiotaomicron* species, along with predicted metabolic pathways across our mouse studies, prompted us to further examine how *B. thetaiotaomicron* affects human intestinal epithelial cells. We focused on the transcriptional landscape and crucial genes and pathways related to IBD wound healing.

### *B. thetaiotaomicron* alters the expression of genes & pathways critical for cell proliferation and migration of cultured intestinal epithelial cells

To evaluate the global impact of *B. thetaiotaomicron* on the transcriptional response of cultured IECs, we performed unbiased RNA-sequencing using **h**uman **p**rimary **c**olonic epithelial **c**ells (HCoEpi) and measured the genome-wide mRNA expression profiles by quantitative deep sequencing of the RNA transcripts (Figure 4). We found marked differences in the mRNA profiles between *B. thetaiotaomicron*-treated and vehicle-treated control cells. Clearly, numerous gene expression patterns in primary HCoEpi cells were different from those in the control (Figure 4A). Among the differentially expressed genes, we observed an increase in genes involved in IEC proliferation (Figure 4B). The heatmap demonstrated the top 50 genes that are associated with IEC proliferation in both human and mouse IECs. We found that *B. thetaiotaom*icron exposure promotes primary HCoEpi cells into a proliferative, mitotically active state, with simultaneous activation of DNA replication, checkpoint, and repair pathways, which are essential for regenerating damaged mucosa. As shown in Figure 4B, the genes involved in cell cycle progression and checkpoints, including *CDK1, CDK2, CDC25A, CCNB1, CCNB2, CHEK1, CHEK2*, and *E2F* transcription factors, were upregulated after *B. thetaiotaomicron* cocultured with primary HCoEpi cells. Similarly, DNA replication licensing genes (*MCM2–MCM7, ORC1, ORC6, and CDC6*) and DNA synthesis regulators (*PCNA, RFC family*) demonstrated elevated expression. Thus, these data suggest that the *B. thetaiotaomicron* stimulates epithelial cells to accelerate entry into S-phase and enhance DNA replication machinery.

**Figure 4.**
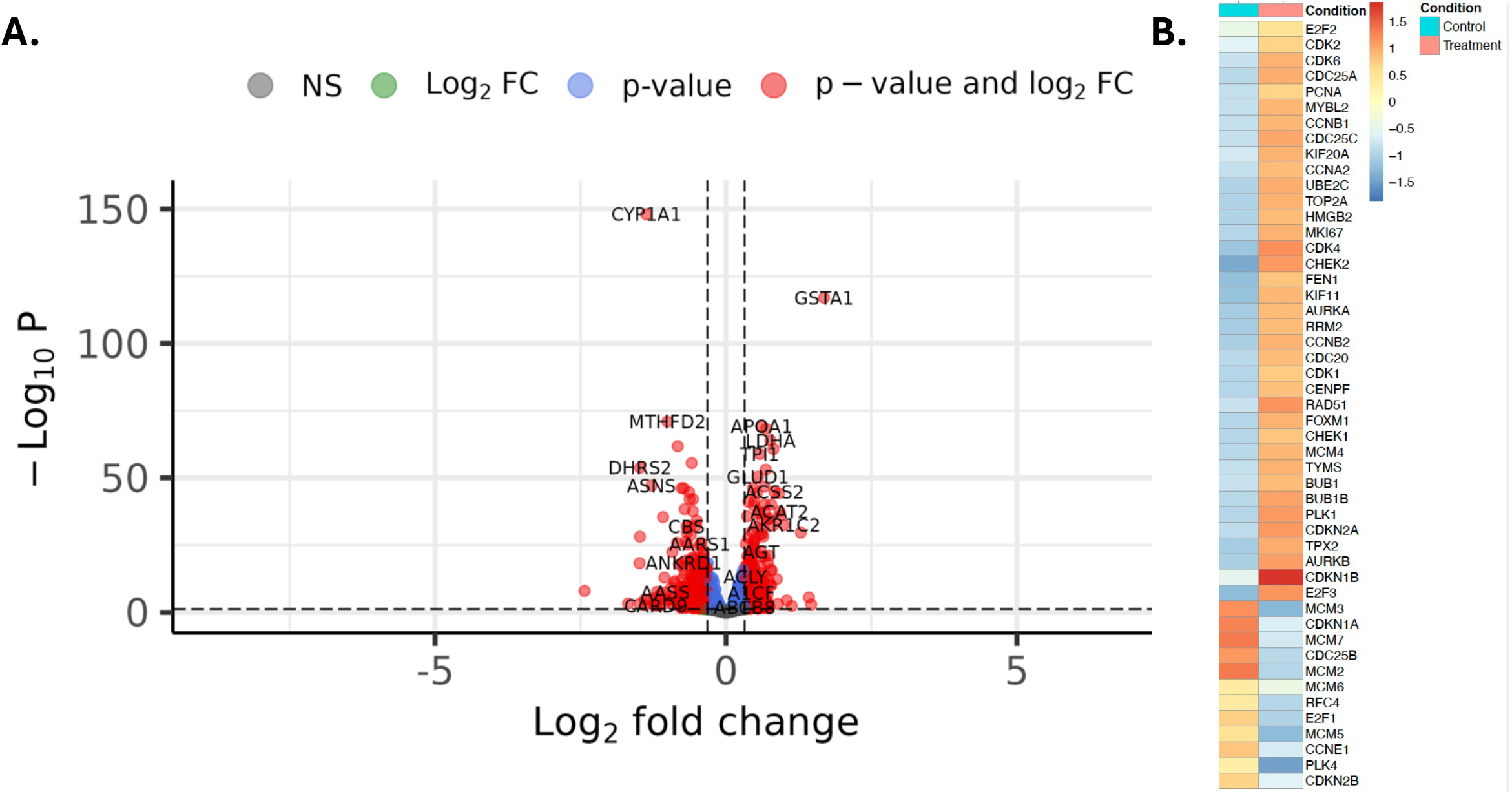
*B. thetaiotamicron* alters the expression of genes and pathways critical for intestinal epithelial proliferation response. (A) Volcano plot showing DEGs stimulated in Caco2 cells after 6 hours of co-incubation with *B. thetaiotamicron*. (B) Heatmap shows the top 50 DEGs involved in proliferation in cultured primary IECs treated with *B. thetaiotamicron* or vehicle. *P < 0.05.

Furthermore, *B. thetaiotaomicron* stimulated upregulation of several spindle assemblies and mitotic regulators, including *PLK1, BUB1B, KIF11, AURKA, and AURKB*. At the same time, stress- and repair-associated genes like *RAD51, TOP2A, and RRM2* were upregulated by the administration of *B. thetaiotaomicron* to the primary HCoEpi cells. Thus, these changes suggest that *B. thetaiotaomicron* may promote epithelial renewal and maintain barrier integrity by enhancing the cell cycle and activating DNA replication, checkpoint, and repair processes. Hence, RNA sequencing of *B. thetaiotaomicron*-treated primary HCoEpi cells revealed increases in many pathways and specific transcripts commonly implicated in IEC proliferation and mucosal repair, including increases in cellular proliferation, differentiation, and migration pathways, providing the impetus for investigation of these pathways in greater depth.

## Methods and materials

### Microbiome Study

DNA was extracted from stool samples (n = 5 mice per group) using a PowerSoil kit from MO BIO Laboratories (Carlsbad, CA)^1^. 16S rRNA genes were PCR-amplified from each sample using a composite forward primer and a reverse primer containing a unique 12-base barcode, designed using the Golay error-correcting scheme, which was used to tag PCR products from respective samples. We used primers for paired-end 16S community sequencing on the Illumina platform using bacteria/archaeal primer 515F/806R. Primers were specific for the V4 region of the 16S rRNA gene. The forward PCR primer sequence contained the sequence for the 5′ Illumina adapter, the forward primer pad, the forward primer linker, and the forward primer sequence. Each reverse PCR primer sequence contained the reverse complement of the 3′ Illumina adapter, the Golay barcode (each sequence contained a different barcode), the reverse primer pad, the reverse primer linker, and the reverse primer. Three independent PCR reactions were performed for each sample, combined, and purified with AMPure magnetic purification beads (Agencourt). The products were quantified, and a master DNA pool was generated from the purified products in equimolar ratios. The pooled products were sequenced using an Illumina MiSeq sequencing platform. Sequences were assigned to OTUs with UPARSE using 97% pairwise identity and were classified taxonomically using the RDP classifier retrained with Greengenes. After chimera removal, the average number of reads per sample was 21,414. A single representative sequence for each OTU was aligned using PyNAST, and a phylogenetic tree was then built using FastTree. The phylogenetic tree was used to compute the UniFrac distances. The PCoA analysis shown is unweighted.

### RNA Extraction

Kit contents were prepared according to instructions. Cells were seeded at 5×10^4^ and grown overnight before treatment with vehicle or *B. theiotamicron* for the appropriate amount of time. On ice, the media was removed from each well, and the cells were washed with chilled D-PBS. 300 μL of RLT+ buffer was added, and cells were scraped off the well using a pipette tip, then transferred into a microcentrifuge tube and vortexed for 45 seconds. 3 μL of β-mercaptoethanol was added to each sample before vortexing for an additional 45 seconds. Cell lysates were transferred to gDNA Eliminator columns and centrifuged at 10,000 rpm for 30 seconds. 600 μL of 70% EtOH was added to the flow-through and mixed by gentle pipetting. Sample was transferred to RNeasy spin columns and centrifuged for 15 seconds at 10,000 rpm, then 700 μL of Buffer RW1 was added to the columns and centrifuged at 10,000 rpm for 15 seconds. Flow-through was discarded and 500 μL of Buffer RPE was added before centrifuging at 10,000 rpm for 15 seconds. An additional 500 μL of Buffer RPE was added, this time centrifuging for 3 minutes at 10,000 rpm. The spin columns were placed into collection tubes, and 40 μL RNase-free water was added to the columns before centrifuging for 1 minute at 10,000 rpm. The flow-through was added back to the column and the centrifugation step repeated. Nanodrop was used to measure RNA concentration.

### qRT-PCR

Reverse transcription was performed according to the protocol in Bio-Rad’s iScript gDNA Clear cDNA Synthesis Kit. qRT-PCR was performed according to the protocol in Bio-Rad’s SsoAdvanced Universal SYBR Green Supermix following instructions for a 10 μL reaction. 2 μL of combined forward and reverse primers were used, along with 2 μL of nuclease-free H_2_O and 1 ng of cDNA template.

### RNA Sequencing

Cells were seeded in 24-well plates at 5×10^4^ and allowed to grow overnight before being treated with PBS or *B. theiotamicron*. RNA was extracted following the protocol above. Quality of the RNA was evaluated using RNA 6000 Nano reagents on the 2100 bioanalyzer. RNA sequencing library preparation was performed utilizing the NEBNext Ultra RNA Library Prep Kit for Illumina by following the manufacturer’s recommendations (NEB). Sequencing libraries were validated on the Agilent 2100 Bioanalyzer System (Agilent Technologies) and quantified using Qubit 2.0 Fluorometer (Invitrogen) as well as by quantitative PCR (Applied Biosystems). The libraries were sequenced on an Illumina sequencer using a 2x150 Paired-End (PE) configuration. Raw sequence data (.bcl files) was converted into fastq files and de-multiplexed. Data were aligned and normalized using STAR aligner. Differentially expressed genes were identified by DEseq2. Gene pathway analysis was performed with the database for annotation, visualization, and integrated discovery (pathway enrichment analyses). We determined statistically significant differences in gene expression at the nominal level of significance (P < 0.05). We also evaluated the effect of Benjamini-Hochberg correction on raw p-values to account for multiple hypothesis testing.

## Discussion

IBD is characterized by dysbiotic gut microbiota, compromised epithelial barrier function, chronic intestinal inflammation, and increased mucosal cytokines and immune cells infiltration. The symbiotic gut microbiota is intricately linked with intestinal health. Many gastrointestinal disorders, including IBD, are immensely impacted by the gut microbiota. However, the microbiota’s nearly 10 million genes and their metabolic functions, which critically influence intestinal health, remained largely unrecognized. Therefore, there is a critical vacuum of knowledge surrounding the specific but mechanistic roles of symbiotic microbial members, their metabolic functions, and the products they generate in the gut. Hence, there has been a remarkable interest in identifying the bacterial genes, pathways, and metabolic products, which may directly impact the functions of the intestinal epithelial cell (IEC) and immune cells of the gut.

Timely and efficient intestinal mucosal wound repair, coinciding with resolution of inflammation, is critical in reestablishing the epithelial barrier and mucosal homeostasis. Resolution of inflammation and repair of the epithelial barrier are mediated by a delicate choreography of epithelium, immune cells, and gut microbiota. Thus, an improved understanding of these biological processes is important in the development of therapeutic strategies designed to promote recovery of epithelial injury in inflammatory states such as IBD.

The gut microbiota beneficially impacts intestinal homeostasis and, by extension, systemic organismal health. There is mounting evidence suggesting that a diverse and balanced gut microbiota benefits maintaining intestinal homeostasis, enterocyte proliferation, differentiation, protection from injury, and epithelial regeneration. Disruption of microbial balance, known as dysbiosis, impairs enterocyte proliferation and migration, leading to compromised epithelial barrier integrity. Altered microbiome composition is frequently associated with changes in bacterial metabolic functions, which can disrupt the biosynthesis of microbial compounds and metabolites. These changes can significantly impact gut mucosal homeostasis, inflammation, and the repair process. However, the detailed mechanisms, changes in commensal microbial metabolic capacity, and their effects on IECs, particularly in altering primary human IEC gene expression, are still not fully understood.

In this project, we studied how the microbiome changes during recovery after mucosal inflammation and epithelial damage caused by DSS. This approach allowed us to model colitis remission, a period characterized by increased repair, including IEC proliferation and migration. We found that after acute colitis, the microbial community structure changes significantly. Alpha diversity decreases, and there are alterations in several taxonomic groups at the phylum, genus, and species levels. During the recovery of injured mucosa, the level of inflammatory Pseudomonodota increases, along with *A. muciniphila* and *B. thetaiotaomicron*. We previously demonstrated that *A. muciniphila* enhances the regeneration of injured gut mucosa by promoting the proliferation and migration of intestinal epithelial cells from the crypts closest to the damaged gut mucosa. Furthermore, in this current study, we found that these changes are associated with decreased Shannon and Simpson indices, indicating that microbial richness and evenness decline one week after DSS is discontinued. Thus, our findings indicate that the dysbiotic gut microbiome does not immediately return to homeostasis or its pre-injury state but undergoes a transitional phase with distinct microbial signatures. Further studies are needed for prolonged follow-up, which would shed light on the trajectory of microbial restoration to either the pre-injury state or a new state that ensures prolonged remission.

Our findings reveal characteristics that both align and contrast the microbiome data from human IBD studies. Multiple studies previously reported a reduced microbial diversity, loss of beneficial Firmicutes (including Bacillota), and an enrichment of *E. coli* and Enterobacteriaceae in IBD patients. Notably, the human intestine affected by IBD often shows a depletion of and *B. thetaiotaomicron*. However, in our recovery-phase model, both species were found to be enriched. This observation aligns with our previous work demonstrating that *A. muciniphila* promotes mucosal restitution and raises the possibility that *B. thetaiotaomicron* may play a pivotal role in epithelial regeneration.

Further study also revealed that *B. thetaiotaomicron* deeply affects the transcriptional landscape of colonic epithelial cells. We found that *B. thetaiotaomicron* treatment stimulated upregulated expression of genes involved in entry into S-phase and mitotic activity, including *CDK1, CDK2, CDC25A, CCNB1, CCNB2, CHEK1, CHEK2*, and members of the *E2F*. Our data also highlighted that DNA replication licensing genes (*MCM2–MCM7, ORC1, ORC6*, and *CDC6*) and replication-associated regulators (*PCNA* and RFC family members) were also elevated, which further supports the notion that *B. thetaiotaomicron* stimulates epithelial cells to accelerate DNA synthesis and division. Moreover, the upregulation of genes involved in stress-response and DNA repair suggests that this B. thetaiotamicron-stimulated proliferative program of IEC is accompanied by activation of checkpoint and repair mechanisms. This dual program suggests that *B. thetaiotaomicron* not only stimulates epithelial renewal but also safeguards genomic integrity.

Overall, these findings elucidate that during recovery from DSS-induced injury, the gut microbiota undergoes a dual shift: compositional changes with expansion of *Bacteroidota* and *B. thetaiotaomicron*, and functional enrichment of metabolic pathways linked to epithelial restitution, stress tolerance, and host–microbe interactions. In addition, transcriptional analysis demonstrated that B. *thetaiotaomicron* stimulates intestinal epithelial transcriptional programs relevant to wound healing and IBD remission. Additionally, the enrichment of pathways linked to differentiation and migration suggests that *B. thetaiotaomicron* may orchestrate multiple layers of the regenerative process. These findings provide important insight into how specific commensal species can influence host repair mechanisms and raise the possibility that *B. thetaiotaomicron* contributes directly to mucosal healing in inflammatory bowel disease and other settings of epithelial injury. This knowledge can be harnessed to promote intestinal homeostasis, prevent relapse, and prolong remission of IBD.

